# A super-resolution protocol to correlate structural underpinnings of fast second-messenger signalling in primary cell types

**DOI:** 10.1101/2020.09.29.319400

**Authors:** Miriam E. Hurley, Thomas M. D. Sheard, Ruth Norman, Hannah M. Kirton, Shihab S. Shah, Eleftheria Pervolaraki, Zhaokang Yang, Nikita Gamper, Ed White, Derek Steele, Izzy Jayasinghe

## Abstract

Nanometre-scale cellular information obtained through super-resolution microscopies are often unaccompanied by functional information, particularly transient and diffusible signals through which life is orchestrated in the nano-micrometre spatial scale. We describe a correlative imaging protocol which allows the ubiquitous intracellular second messenger, calcium (Ca^2+^), to be directly visualised against nanoscale patterns of the ryanodine receptor (RyR) Ca^2+^ channels which give rise to these Ca^2+^ signals in wildtype primary cells. This was achieved by combining total internal reflection fluorescence (TIRF) imaging of the elementary Ca^2+^ signals, with the subsequent DNA-PAINT imaging of the RyRs. We report a straightforward image analysis protocol of feature extraction and image alignment between correlative datasets and demonstrate how such data can be used to visually identify the ensembles of Ca^2+^ channels that are locally activated during the genesis of cytoplasmic Ca^2+^ signals.

## 1. Introduction

Single molecule localisation microscopy (SMLM) techniques have grown into some of the most useful fluorescence imaging techniques available to cell biologists. Techniques like photo-activated localisation microscopy (PALM) and direct stochastic optical reconstruction microscopy (dSTORM) have offered unique visual insights into protein organisation (1–4), membrane-bound organelles (1, 5, 6), cytoskeletons (7, 8), signalling complexes (9, 10) and nano/microdomains (11–13). Key to their increasing popularity is the versatile and inexpensive strategies to visualising biological structures at the true molecular scale (i.e. at the spatial scale of ~ 10-50 nm) with high spatial precision and high molecular specificity. Protocols like DNA-PAINT (14) have extended the achievable spatial resolution to 1-10 nm, and have enabled image-based omics such as target counting (15, 16), proximity mapping (17) and multiplexed super-resolution imaging (14, 18).

A significant bottleneck in the utility of SMLMs when studying dynamic biology, such as intracellular signalling, is that it is inherently time-consuming. Super-resolution images from these methods are typically reconstructed stochastically based upon the photoswitching of markers which are localised at the target structure. Generally, these imaging protocols can only resolve living sub-cellular structures with a time-resolution of seconds (19, 20) to minutes (21), without the aid of accelerating algorithms. Maintaining cell health whilst delivering the targeted fluorescent probes across the plasmalemma is also challenging, As is managing the photodamage imparted by the high excitation powers required for photoswitching, and the altered oxygen availability in the presence of switching buffers (22, 23).

Genetically-encoded markers, particularly photoswitchable fluorescent proteins (FPs) like green FP (GFP) offer a useful workaround. For biomedically-interesting samples however this may still be limiting. Cell, tissue or organism types which are commonly associated with specific disease conditions or vital physiology are generally not available with natively expressed FPs. For example, animal models could be used to express ion channels (24) which are critical to the development and homeostasis of all mammalian excitable tissue types, such as the ryanodine receptor type-2 (RyR2). However, for many groups studying the biophysics of proteins or cellular structures which are critical for normal cell function, these animal models are beyond their technical, collaborative and/or financial means. *In vitro* gene transfection of primary cells with FP-fusion protein constructs are an alternative (25, 26); however marker sparsity, depending on transfection efficiency and native protein turnover rates, are a major drawback. Cell types such as muscle also undergo severe maladaptive remodelling *ex vivo* (27) which limits the utility of gene transfection as a route of delivering the protein markers for super-resolution imaging. For primary cell-types which must be dissected and/or isolated acutely prior to imaging, a correlative imaging protocol which includes sequential live-cell and correlative SMLM is therefore ideal.

Junctional membrane complexes (JMCs) are a type of intracellular structure which are challenging to study in transgenic animals or in vitro models. They are sub-micron-scale intracellular signalling hubs which are responsible for fast intracellular calcium (Ca^2+^) signalling. Elementary Ca^2+^ signals, known broadly as ‘Ca^2+^ sparks’(28, 29) are produced from individual or groups of JMCs. They are one of the most ubiquitous second messenger mechanisms, responsible for a variety of vital physiological processes. In the heart, Ca^2+^ sparks form the primary trigger for the contraction of myocytes which power the blood circulation. Additionally, they are also the basis of skeletal and smooth muscle contraction, Ca^2+^ dependent release of neurotransmitters and pain-related neuropeptides in neurons, fertilization of vertebrate eggs and the secretion of hormones from glandular epithelial cells (see Cheng & Lederer (30) for an overview). RyRs residing within JMCs are the most common source of Ca^2+^ sparks; however other intracellular channels such as inositol triphosphate receptor (IP3R) are also implicated in the genesis of these elementary signals and are thought to be at the centre of many pathologies such as cardiac arrhythmias and myopathies. A need for correlative imaging springs from the limited understanding of how the *native* geometry of organisation of these ion channels determines the spatial patterns of the Ca^2+^ sparks present across a variety of cell types.

Ca^2+^ sparks can be commonly imaged using AM-loadable, high-affinity fluorescent calcium indicators. The principal Ca^2+^ channels and transporters responsible for this signalling are organised in highly intimate clusters within the JMCs (11, 31). Our previous work has shown that discerning individual channels within those clusters requires an optical resolution no worse than ~ 15 nm (4, 11). Therefore, an analysis which looks to examine the relationship between the co-clustering patterns or cluster sizes of these Ca^2+^ channels requires a true-molecular scale imaging method such as DNA-PAINT, which is capable of mapping and quantifying marker patterns at a resolution < 10 nm.

Correlative microscopy protocols have occupied the area of experimental biophysics over many decades now. Currently, one popular implementation is correlative light-electron microscopy (CLEM), which combines the ångström-scale resolution of electron microscopies (EM) with the molecular specificity of fluorescence markers (31). Other combinations of techniques that have been demonstrated recently include super-resolution/atomic force correlative microscopy (32–35) and optical multi-modality (36–38) correlative protocols. By a considerable margin, most of these protocols exploit the different resolving powers, spatial dimensionalities and/or specificity of the combined techniques to map structures in order to reconcile cellular topologies and limitations within the individual imaging modalities. However, the correlative approach remains under-utilised for mapping dynamic (functional) features such as ionic signals or membrane potential against the nanoscale structures which underpin them.

This paper outlines a correlative optical imaging pipeline that we developed with the aim of reconciling the dynamic Ca^2+^ signalling (diffraction-limited) patterns within living primary cells with the underlying channel organisation patterns. To this end, we demonstrate the alignment of sub-plasmalemmal patterns of fast and spontaneous cytoplasmic Ca^2+^ signals, acquired at diffraction-limited resolution, to super-resolution images of RyR Ca^2+^ channels in rat cardiomyocytes (principle illustrated in Fig 1).

**Figure 1:**
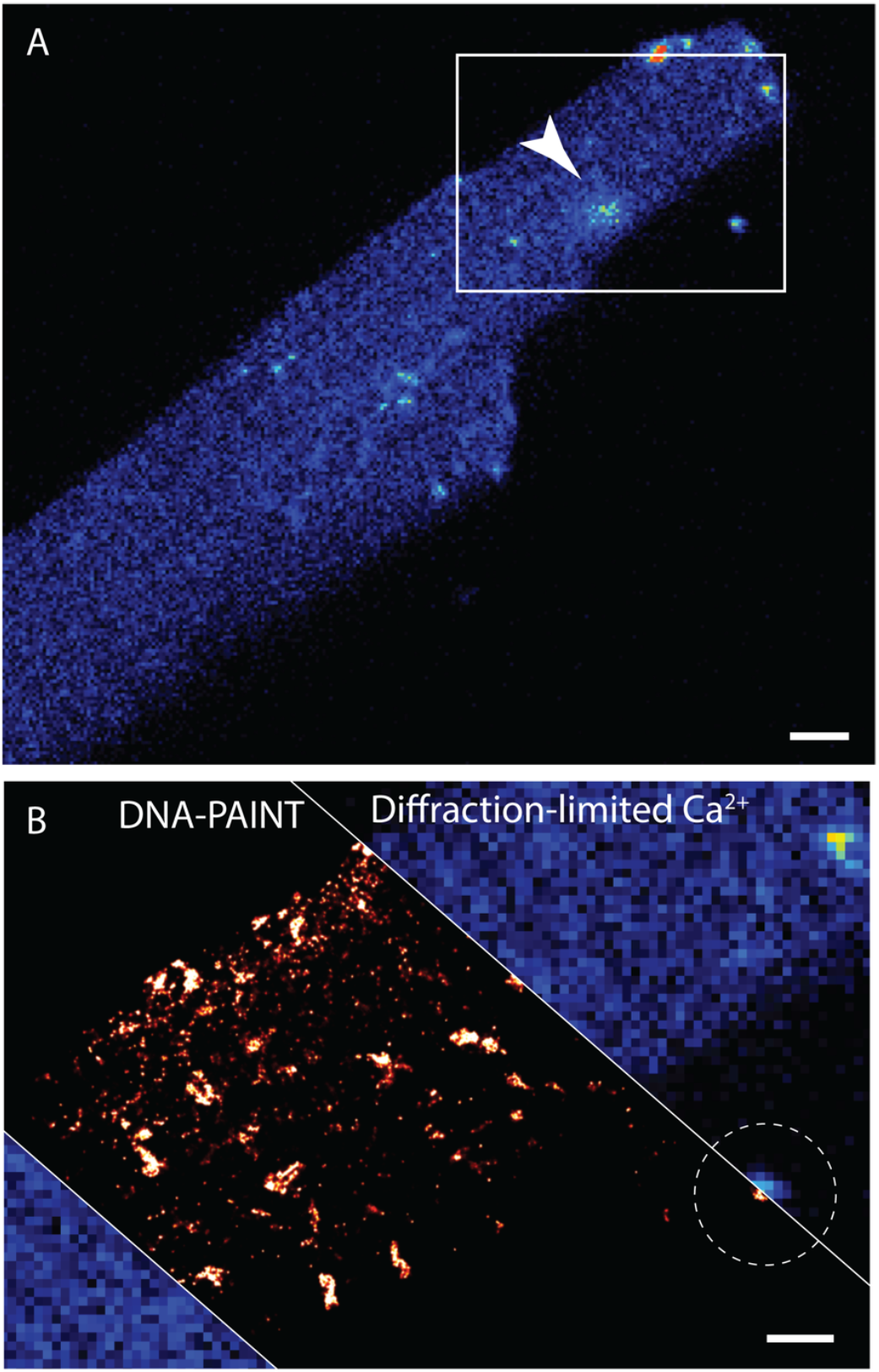
The principle of correlative diffraction Ca^2+^ and super-resolution Ca^2+^ channel mapping. The method relies on initially imaging fast, two-dimensional, diffraction-limited time series of Ca^2+^-sensitive fluorescence of cytoplasmic Fluo-4 AM dye recorded within a living cell. (A) A single frame of a rat ventricular myocyte exhibiting a single Ca^2+^ spark (arrowhead). (B) Magnified view of the window indicated in panel A, overlaid (diagonal strip) with a super-resolution DNA-PAINT image of the ryanodine receptor clusters acquired subsequently. The correlative images are registered with the alignment of either fluorescent fiducial markers (dashed line) and/or the alignment of the cell outline. Scale bars, A: 4 μm, B: 1 μm.

## 2. Microscope set up

Experiments were performed on a Nikon TE2000 (Nikon; Japan) microscope with an after-market Physik Instrumente stage (Model: M-545) containing a circular petri-dish inset (M-545.PD3). The microscope consisted of a custom light path which followed the schematic shown in Figure 2. Total internal reflection fluorescence (TIRF) illumination was used for both the DNA-PAINT and the Ca^2+^ imaging experiments in order to ensure that the images were recorded from within an evanescent field at the surface of the cell. All imaged cells were attached to the #1.5H coverslip bottom of a glass dish (Ibidi, UK).

**Figure 2:**
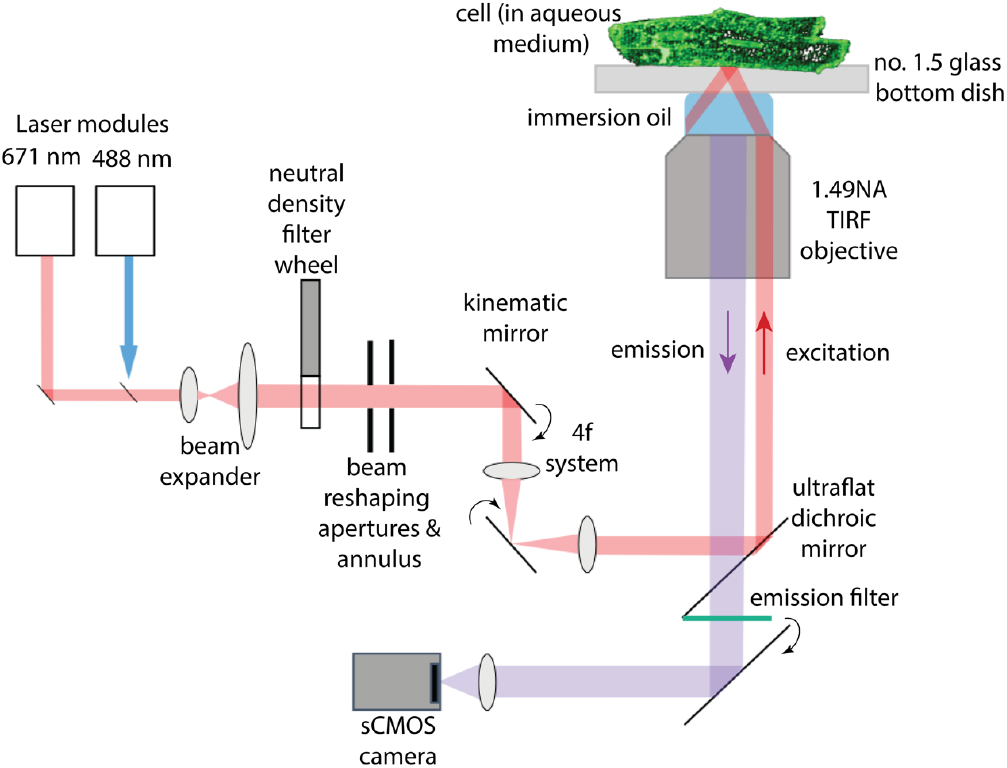
Light path of the TIRF microscope used for performing the Ca^2+^ imaging and DNA-PAINT.

For DNA-PAINT, fluorescent markers were excited using a 642 nm CW diode laser with a power output of 800 mW (Viasho, China). A 488 nm Cobolt Jive DPSS laser (Cobolt AB, Solna) of equivalent power was used for the Ca^2+^ imaging. A computer-controlled motorised neutral density filter wheel (Thorlabs, Germany; FW102C) was used as the primary method of regulating the excitation laser power. The laser beam was positioned onto the periphery of the rear aperture of a Nikon 1.49NA TIRF objective using a 4f beam steering system in such a manner that achieves a collimated super-critical excitation beam at the front of the lens. This ensured that the TIRF evanescent field was positioned as a thin horizontal light sheet at the surface of the cell. Using one of the fine grid regions of a PSFcheck multi-emission reference slide (PSFcheck, Exeter), the 488 nm and 642 nm TIRF illumination spots were aligned in the x-y plane in order to ensure that the Ca^2+^ and the DNA-PAINT images were acquired broadly within the same region of the field of view.

Ultraflat dichoric mirrors (Chroma T495lpxr 1 mm dichroic for 488 nm excitation and Chroma T685lpxr 1 mm dichroic for 642 nm excitation) were used to split the emission light from the excitation path. Emission filters ET525/50m and ET720/60m (Chroma) were required for the two spectral channels of imaging, respectively. Emitted light was recorded onto a Zyla 5.5 USB scientific CMOS camera (sCMOS; Andor, Belfast).

Imperfections in the flatness and/or alignment between the differential emission filters and beamsplitters as well as misalignment in the two light paths of the 488 nm and 642 nm lasers can introduce local misalignments *between* the two images. A one-off distortion-correction vector map was generated by imaging a sparse distribution of multi-spectral 100-nm Tetraspeck beads (Thermo Fisher) with the two respective imaging channels. This correction vector map was used for locally adjusting the coordinates of the Ca^2+^ sparks relative to the DNA-PAINT channel.

Image acquisition was done on a Lenovo Thinktank workstation using an Intel i7 quad-core processor, 32 Gb of DDR3 memory and a 2 Tb solid-state drive running the open-sources Python Microscopy Environment (PyME) software (freely-available for download via www.python-microscopy.org). The PyME ‘Aquire’ image acquisition interface included a live display of the camera’s acquisition at a customizable frame-rate. This was particularly critical for acquiring reference images of the sample between steps, for orientating the sample and for positioning the cells’ region of interest within the illumination TIRF field.

## 3. Cell preparation

The protocol below was applied to ventricular cardiomyocytes from adult male Wistar rats as detailed previously (11). All animal experiments were conducted according to the UK Animals (Scientific Procedures) Act of 1986 under the EU Directive 2010/63/EU with UK Home Office and local ethical approval.

### 3.1. Cell isolation

To track a cell across the duration of the protocol, 500 μm square gridded dishes with #1.5H glass coverslip (Ibidi, UK) were utilised (Figure 3A).

**Figure 3:**
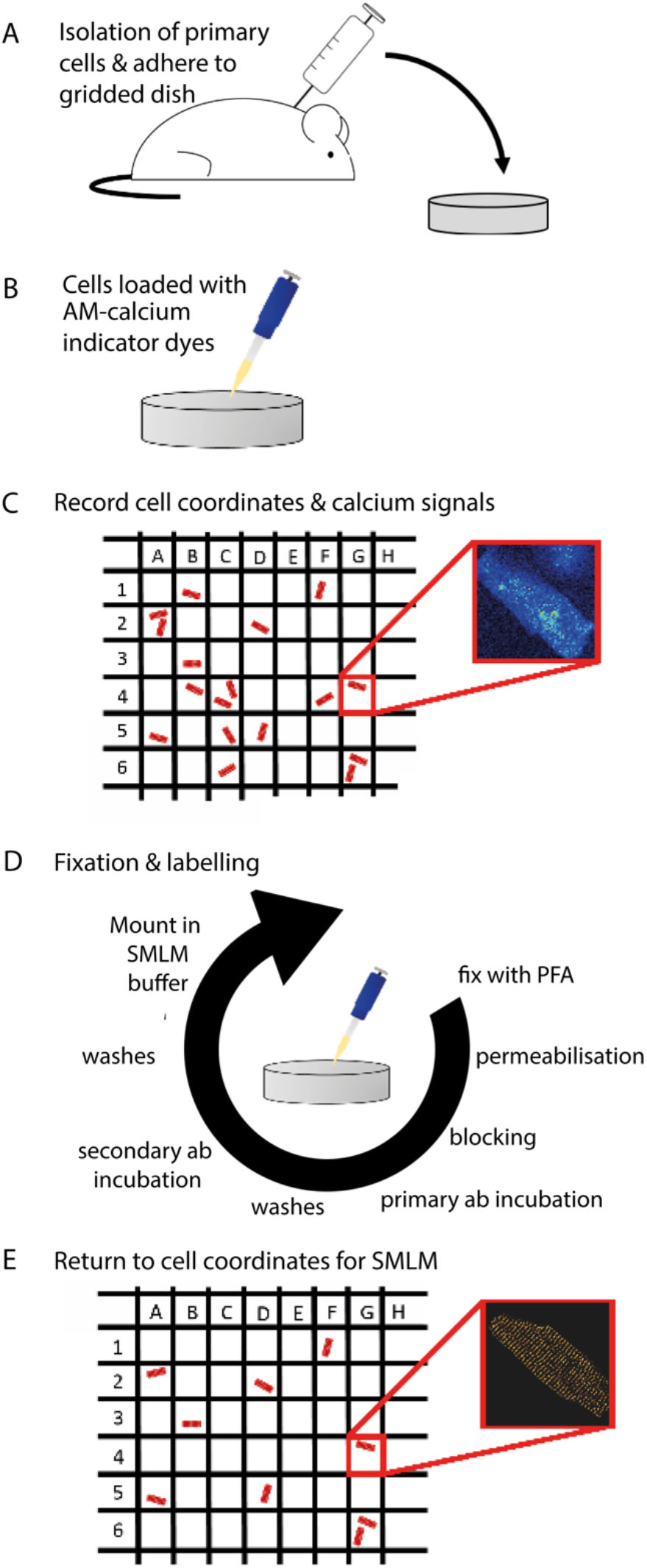
The experimental pipeline of correlative Ca^2+^ and SMLM in primary cells. The key steps include (A) Extracting and isolating living cell type(s) of interest from the organism of interest and adhering or culturing them within the gridded dishes; (B) AM-loading of the calcium indicator dye and de-esterification; (C) recording Ca^2+^-sensitive indicator dye fluorescence from cells and the specific grid coordinates noted; (D) fixation and immune-labelling towards SMLM; (E) return to the coordinates of a cell whose Ca^2+^ signals were recorded previously.

To ensure cell attachment, dishes were coated overnight with 11.9 μg/mL laminin for cardiomyocytes.

Ventricular cardiomyocytes from adult male Wistar rats were obtained from enzymatic isolation upon the Langendorff apparatus (detailed previously in the supplementary section of Jayasinghe & Clowsley et al (11)). Briefly, the whole heart was cannulated at the aorta. To clear the coronary circulation, isolation solution with added 0.75 mM CaCl_2_ was perfused at 7 mL/min. Isolation solution at pH 7.4 constituted of (in mM) 130 NaCl, 1.4 MgCl_2_, 5.4 KCl, 0.4 NaH_2_PO_4_, 5 HEPES, 10 Glucose, 20 Taurine, 10 Creatine. Once cleared, isolation solution with added 0.1 mM EGTA (Sigma-Aldrich, USA) was perfused for 4 minutes. Enzymatic digestion of the whole heart followed. Isolation solution was perfused for 7 minutes with 1 mg/mL Collagenase Type II (Worthington Biochemicals, USA) and 1.8 mg/mL protease (Sigma-Aldrich, USA). Atria and vasculature were dissected away. The remaining ventricles were triturated to liberate isolated cardiomyocytes into a suspension which was used in the experiment without further delay.

## 4. Live cell (Ca^2+^) imaging

Prior to live-cell Ca^2+^ imaging, cells were exposed to their cell-specific Tyrode’s solution. This constituted of (in mM) 140 NaCl, 4 MgCl_2_, 1 KCl, 10 HEPES, 10 Glucose, 5 CaCl_2_ at pH 7.4 for cardiomyocytes. Pre-exposure to 5 mM CaCl_2_ Tyrode’s solution identified those cells which were tolerant to high extracellular calcium, increased the propensity of a cell to produce spontaneous Ca^2+^ spark activity, and enabled cells to be selected for quiescence (mechanically relaxed state) during the live cell phase of the experiment. One could also use ‘uncoupling’ drugs such as 2,3-butanedione monoxime (BDM) which would limit contractile motion of myocytes, however we did not opt for it because of its known interactions with intrinsic Ca^2+^ signalling and cellular excitability (39).

### 4.1. Loading of cells with calcium indicator dye

Cells were loaded with 5 μM Fluo-4 AM (ThermoFisher Scientific, UK) for 15 minutes at room temperature (RT; Figure 3B). According to Steele & Steele (40), cells were washed twice with their respective Tyrode’s solution to remove excess Ca^2+^ indicator dye prior to a 30 minute de-esterification stage at 4°C. A third wash was performed.

After dye loading, cardiomyocytes were allowed to self-adhere to a glass-bottom imaging dish. Cardiomyocytes demonstrated better Ca^2+^ indicator dye loading when it was performed prior to their addition to the imaging dishes. In addition, TetraSpeck microspheres (0.1 μm sized; ThermoFisher Scientific, UK) were added to act as fiducial markers. Cells were incubated for 90 minutes at RT prior to imaging.

### 4.2. Ca^2+^ spark imaging with Total internal reflection microscopy

According to the grid lines, dishes were clamped securely on a the cicular Physik Instrumente stage inset on the TIRF microscope system. Using brightfield illumination, the dish was oriented in such a manner that the grid lines were perfectly aligned with the edges of the live image of the camera. This could be continuously monitored using the ‘live view’ displayed on the PyME ‘Acquire’ software interface. The microscope was switched onto 488 nm TIRF illumination and cells which form a substantial footprint with the dish were identified. At 100 ms/frame, and a laser illumination power of ~1×10^6^ W/cm^2^, spontaneous Ca^2+^ spark activity within frames which ideally contain both the cell’s footprint of attachment and multi-spectral fluorescent bead fiducials were recorded onto 12-bit HDF time series image files (.h5 which can be opened via PyME). A still image was acquired under brightfield illumination, following each Ca^2+^ spark series which provided a note of the cell’s orientation within the image and the gross coordinates within the grid of the dish (schematically shown in Figure 3C).

## 5. Fixation, Immunofluorescence staining & DNA-PAINT

### 5.1. Fixation

Immediately after Ca^2+^ imaging, cells were fixed *in situ* with 2% paraformaldehyde (Sigma-Aldrich; w/v in phosphate buffered saline; PBS) for 10 minutes at room temperature (RT). Fixative was removed with PBS over 4× 10-minute washes at RT.

### 5.2. Immunolabelling

Cells were permeabilised for 10 minutes at RT with 0.1% Triton X-100. This was followed by the application of a PBS based blocking buffer, for 60 minutes at RT, which contained 10% normal goat serum.

1. To label a specific protein of interest, cells were incubated overnight at 4°C with an anti-RyR2 primary antibody (1:250; Sigma-Aldrich; Cat# HPA020028). The antibody was diluted to its required working concentration using incubation solution. Incubation solution consisted of (w/v or v/v); 0.05% NaN_3_, 2% BSA, 2% NGS and 0.05% Triton X-100 dissolved in PBS.
2. Cells were washed 3 times for 20 minutes each in PBS. Following this, a secondary antibody conjugated to a DNA-PAINT ‘docking strand’ (sequence: C6 dT-amine - 5’ TTA TAC ATC TA 3’; custom made via Integrated DNA Technologies) was applied, again diluted at 1:100 in incubation solution, for 2 hours at RT. These secondary antibodies (unlabelled goat anti-rabbit-IgG; Jackson Immuno Research) were conjugated to the ‘docking strand’ using the protocol outlined previously (11).

## 6. DNA-PAINT image acquisition

The ‘imager’ strands (sequence: 5’CTA GAT GTA T3’-Atto655 custom made via Eurofins), complementary by sequence to the ‘docking strands’, were diluted at a concentration of 1.2 nM in ‘Buffer C’ (PBS with final 500 mM NaCl; pH 8.0) previously described by Jungmann et al. (14). The dish was placed on the microscope stage and the cell was re-positioned within the imaging field using the grid coordinates recorded in the Ca^2+^ imaging step. As in the previous imaging step, the chamber was rotated within the stage inset until the grid-lines were parallel to the edges of the ‘live-view’ which is available in the PyME ‘Acquire’ interface. In our observation, this ensured that any angular offsets between the Ca^2+^ and the DNA-PAINT data were eliminated.

During image acquisition, an Atto 655-conjugated oligonucleotide ‘imager’ strand (1.2 nM) was added to Buffer C. Under TIRF illumination, the thermally-driven reversible binding of the imager strands to the docking strands located within the evanescent field gives rise to bright events within the cell’s ‘footprint’ of attachment, as described previously (11). The stage was re-positioned in the x-y plane such that the visible footprint of attachment containing the DNA-PAINT events was aligned approximately with the area of the field of view where the corresponding Ca^2+^ sequence was recorded (i.e. well within the illumination field). Time series of the DNA-PAINT events were acquired at 100 ms/frame integration time using the same software, and saved in the same image format. The laser illumination used was within the range of 1 – 5 × 106 W/cm^2^. For a typical DNA-PAINT image, 20,000-40,000 frames were acquired.

## 7. Image processing

### 7.1. Ca^2+^ image data

The Ca^2+^ indicator image series contained fluorescent events which were either spontaneously-occurring Ca^2+^ sparks localized to sub-cellular regions and lasting 1-2 frames or, intermittently, larger and more global events such as ‘Ca^2+^ waves’. These time series were used in two distinct pipelines of analysis. Firstly, ~ 10 consecutive frames of the series were used to develop an averaged (low-noise) image which was taken as a reference image of the cell’s boundaries. The cytoplasmic [Ca^2+^] at the resting state provided a low intensity, diffraction-limited image clearly demarcating the cell’s boundaries (Figure 4A). This averaged image provided a low-noise reference image for determining the alignment of the Ca^2+^ images, and therefore the Ca^2+^ spark positions, in relation to the DNA-PAINT images of the RyR organisation (see below).

**Figure 4:**
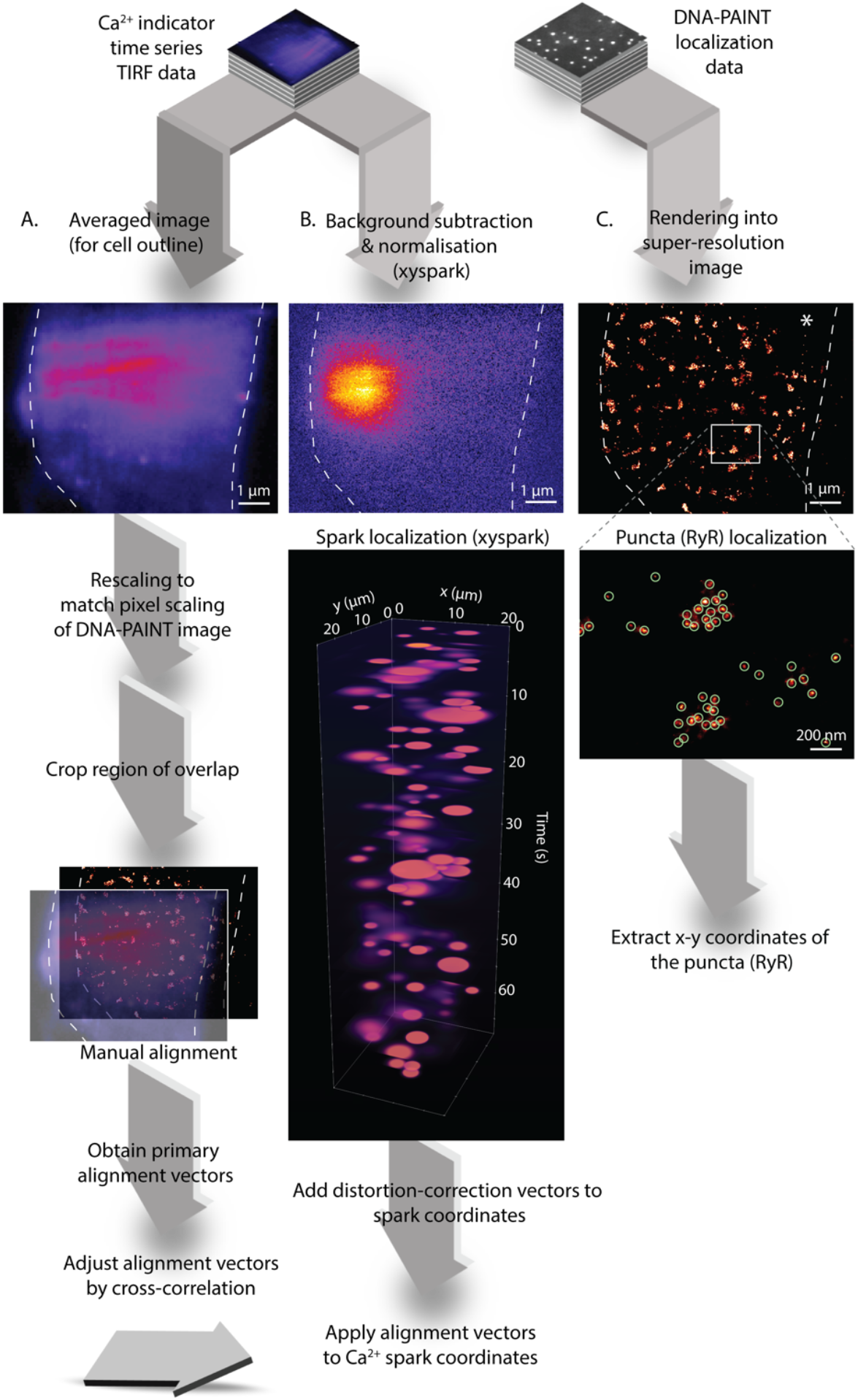
Feature alignment roadmap. (A) The time series of TIRF Ca^2+^ images are used for constructing a low-noise averaged image of the cell geometry which is used as a proxy for determining the alignment vectors for registering Ca^2+^ data against the DNA-PAINT data. (B) The same Ca^2+^ image series are used for Ca^2+^ spark localisation via ‘xyspark’ which involves a background subtraction and spark localization through time to determine their positions, amplitudes and variable sizes. These coordinates are then shifted using the alignment vectors. (C) The DNA-PAINT localization data are used for rendering a super-resolution image of the RyR pattern; these images are then used for localizing and discretizing the punctate RyR densities.

Secondly, the .h5 data volumes were extracted into .tiff stacks and used for detecting Ca^2+^ sparks and localizing them using the xyspark plug-in (40) in FIJI (Figure 4B). Briefly, xyspark required the user identification of frame(s) devoid of Ca^2+^ spark or wave activity which are then used as a denominating frame for a background removal. It then detects the Ca^2+^ sparks (e.g. top panel of Figure 4B), and performs a localisation protocol which fits a 2D-Gaussian shape to the sparks throughout the entire time-series (lower panel of Figure 4B). The localization outputs of the plug-in include the x,y and time coordinates of the sparks, their full-width at half maximum (FWHM) estimated by the Gaussian fit, coefficient of determination *R*^*2*^, amplitude estimated as *F*/*F*_*0*_, where *F*_*0*_ was the estimate of the baseline level of the Ca^2+^ indicator fluorescence in a local cytoplasmic region and *F* was the fluorescence intensity value at the peak of the spark, estimated by the Gaussian fit. In general usage, *F*/*F*_*0*_ is an estimation of the height of a Ca^2+^ signal as normalised to the local baseline fluorescence. To select localization parameters of Ca^2+^ sparks which were in-plane, elementary and not spurious, only the sparks with 1.0 μm ≤ FWHM ≤ 6.0 μm **and** an *R*^*2*^ value ≤ 0.5 were retained for further analysis.

### 7.2. DNA-PAINT image data

DNA-PAINT primary data (image series) were recorded onto .h5 datasets. In addition, a live analysis was performed to localise individual marker events using the dh5view interface of PyME, as described previously (11). Localized point coordinates along with the localization parameters were saved into a more compact format, .h5r, which could be retrieved later. The point coordinates could later be visualised offline using an additional interface of PyME called VisGUI. The DNA-PAINT localization data were rendered into a super-resolution 32-bit greyscale image at a pixel size no larger than 5 nm/px using a rendering algorithm described previously by Baddeley et al. (41). In the rendered images (e.g. Figure 4C), the local intensity reflected the local event density weighted by the localization precision.

The rendered DNA-PAINT image data exhibited a distinct punctate pattern which was characteristic of the RyR cluster patterns observed near the cardiac muscle cell plasmalemma (previously with DNA-PAINT (11) and expansion microscopy (4)); each punctum was equivalent to individual RyRs). A localization protocol used previously (see supplementary section of Jayasinghe & Clowsley et al. (11)) was used to determine the x-y coordinates of individual RyR puncta (detections shown as green circles in lower panel of Figure 4C). This discretisation of RyRs was a critical step in performing spatial statistics following the alignment of the correlative data.

### 7.3. Image alignment

Given that the Ca^2+^ image data and the DNA-PAINT data were not acquired neither simultaneously or immediately in series, the two sets of correlative image data required careful alignment. This registration aimed to remove in-plane spatial offsets in re-seating the sample following cell fixation and immunostaining.

Using custom-written code implemented in IDL v 8.0 (L3HARRIS), the averaged Ca^2+^ image was rescaled to match the pixel scale of the greyscale DNA-PAINT data (typically 5 nm/px). Both the DNA-PAINT image and the averaged Ca^2+^ image were cropped in order to remove non-overlapping regions of the two respective images and to give both images the same dimensions. Using the RGB display windows of IDL and custom-written image shifting commands, the averaged Ca^2+^ image was iteratively shifted into position by a human user such that the cell outlines (and any fiduciary marker beads present nearby) were in approximate (coarse) alignment. These x and y shift vectors were recorded as the ‘primary alignment vectors’.

An additional level of automated fine alignment was performed where the coarsely aligned image pairs were subjected to a 2D cross-correlation. Deviations of the peak correlation value from the centre of the 2D result was used to calculate a fine adjustment to the primary alignment vectors. In scenarios where the fine alignment vectors exceeded 300 nm, the alignment reverted to the primary alignment vectors.

An example dataset and the IDL code used for the image alignment steps is available on the online GitHub repository https://github.com/ijayas/imagealigning including a step-by-step guide to executing this code.

#### Aligning the Ca^2+^ spark coordinates relative to the DNA-PAINT image

The x and y coordinates of the Ca^2+^ sparks (in the uncropped version of the data) were individually shifted using the distortion correction vector field used for estimating the in-plane shifts between primary 488 nm and 642 nm channels (see the Microscope setup section). The widths of the bottom and left margins cropped out from the averaged Ca^2+^ image early in the image alignment step, were subtracted from x and y coordinates of the Ca^2+^ sparks respectively. All of the sparks which remained within the cropped window were then shifted uniformly by the primary alignment vectors, in order to bring them in alignment with the rendered DNA-PAINT image. These Ca^2+^ spark coordinates were now aligned with the coordinates of their true underlying RyRs.

### 7.4. Local interrogation of the RyR patterns & spark-based spatial statistics

The spatial alignment of the Ca^2+^ image data with the true molecular scale RyR patterns, visualised with correlative DNA-PAINT (Figure 5A) now enabled us to examine the features or RyR organisation that are local to the Ca^2+^ sparks. The two sets of coordinates (the localized Ca^2+^ sparks along with their localization parameters, and the RyR puncta positions) offered an opportunity to directly examine their spatial relationship. Ca^2+^ sparks are large, diffusive events (visually represented in Figure 5B). Their large size owed to multiple factors including the diffusion of Ca^2+^ released from localised groups of RyRs (shown with a two-dimensional visualisation in Figure 5C), the diffusion of the de-esterified Fluo-4 (AM) dye within the cytoplasm and the buffering of Ca^2+^ within the cytoplasm. Working within these uncertainties, the FWHM of the spark estimated by xyspark (indicated by the black dashed lines in Figure 5B, or solid-lined circle in Figure 5C) offered a spatial sample of the microenvironment within which RyRs are likely to be activated in order to produce the spark (white dashed lines). One analysis enabled by this sampling strategy was a straightforward counting of the RyR channels underpinning the spark. The integral of the Ca^2+^ signal associated with the spark, defined as ‘spark mass’ (43), was calculated as the product of the spark amplitude (*F*/*F*_*0*_), FWHM^3^ and conversion factor 1.206, and was one of the default output parameters of the xy spark software.

**Figure 5.**
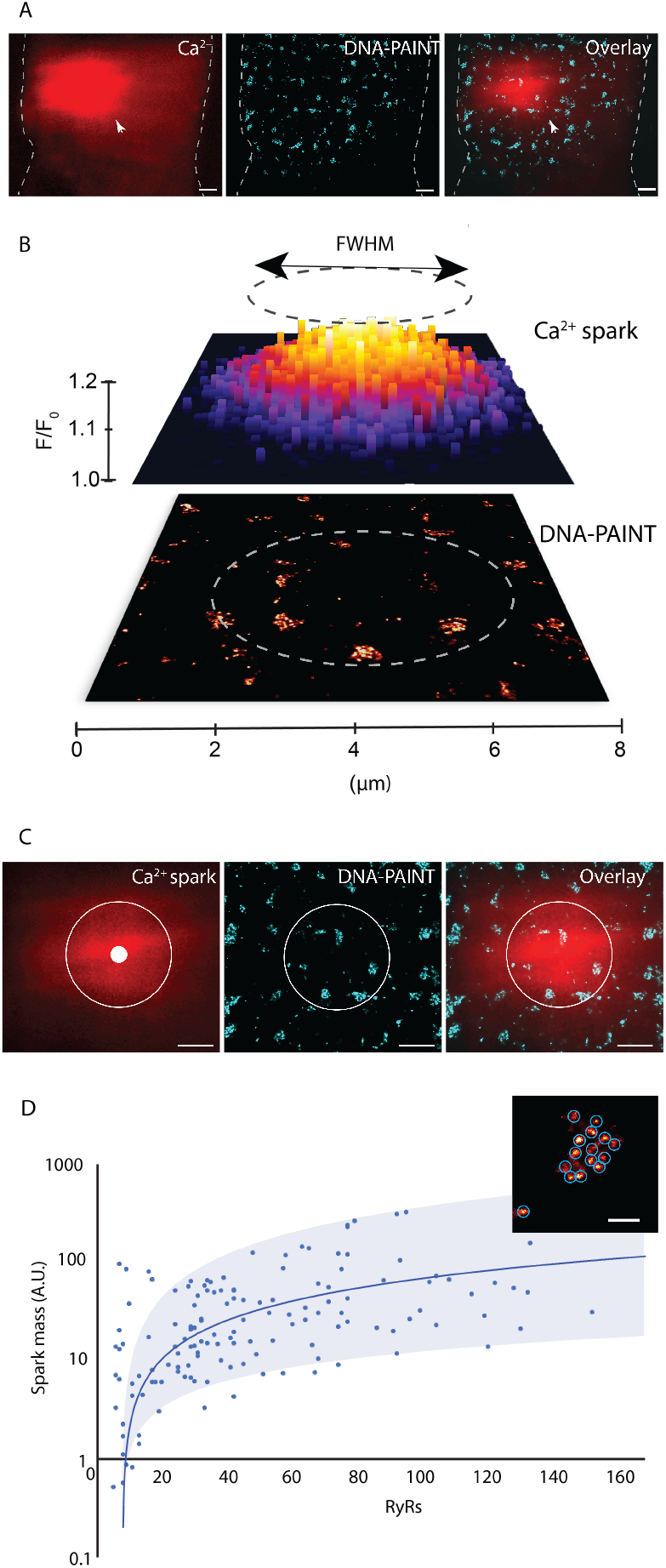
Local interrogation of the RyR channels underlying the Ca^2+^ spark. (A) Alignment of one of Ca^2+^ image frames containing the cell overview (red) indicating one calcium spark (arrowhead) and the corresponding RyR DNA-PAINT image is shown (overlay on right). Dashed line illustrates cell outline. (B) For the local interrogation, the full-width at half-maximum (FWHM; circle with black dashed line) of the localised Ca^2+^ spark was used as a window of confidence (circle with white dashed line) to examine the the DNA-PAINT image (bottom image) for RyRs which were likely to have been local, and therefore likely to have contributed Ca^2+^ release to the spark. (C) A two-dimensional view of a Ca^2+^ spark (red), its centroid detected using the xyspark software (white dot) and the interrogation window of confidence (circle with solid white line). The RyRs resolved with DNA-PAINT (cyan) inside this window of confidence (centre), also shown in overlay (right). (D) RyR counting (as shown in inset) under independent Ca^2+^ sparks indicated the positive relationship between the integral of the Ca^2+^ released (‘spark mass’) and the total number of RyRs Scale bars: A: 1 μ;m, C: 2 μ;m & D: 100 nm.

Figure 5D shows a scattergram of this spark mass against the number of RyR puncta illustrating the typical numbers of RyRs in cardiomyocyte sub-plasmalemmal nanodomains which physically underlie the Ca^2+^ sparks. It revealed that ~ 95% of the sparks arise within local regions which contain between 8 and 105 RyRs. The points cloud in the scattergram showed a significant deviation on either side of a predicted trend-line (shaded regions indicate ~ 90% confidence interval), illustrating that the number of available RyR channels is only a loose predictor of the local Ca^2+^ spark signal.

The IDL script required for sampling and correlating the RyR channels based on the Ca^2+^ sparks is included in the GitHub repository https://github.com/ijayas/imagealigning alongside a step-by-step guide to executing this code and a dataset.

The key image processing steps are summarised in a step-by-step sequence in Table 1, along with the rationale, software used and the tolerances relevant for a correlative analysis similar to that presented in this paper.

**Table 1:** Step-by-step guide to the image processing pipeline, the rationale and the software tools required.

|  | Analysis step | Rationale | Software and approach used | Tolerances |
| --- | --- | --- | --- | --- |
| 1. | Estimation of an averaged (low noise) 2D image of the Ca<sup>2+</sup> image series | A low noise image from the Ca<sup>2+</sup> channel is required for obtaining a reliable impression of the cell outline, useful in the manual image alignment. | FIJI: Take an average intensity projection between consecutive frames devoid of Ca<sup>2+</sup> sparks.<br>The image is used in step 5 of this table for manual image alignment. | The 2D area of the time series may require cropping to ensure only the regions of the cell and surrounding with correlative features are included. This helps conserve computer memory as the images are upsampled in step 5. |
| 2. | Ca<sup>2+</sup> Spark detection | For a spark-by-spark interrogation of the local RyR patterns, x-y coordinates of the centroid and the FWHM are required. | ‘xyspark’ plugin by Steele & Steele (41), implemented in FIJI. The plugin will instruct the user on the required steps towards automatically estimating and subtract background.<br>The final output of the fitting and localisation parameters of the Ca<sup>2+</sup> sparks can be saved as a .txt file. | No tolerance required for the detection. Only sparks with $1.0 \mu\text{m} \leq \text{FWHM} \leq 6.0 \mu\text{m}$ and an $R^2$ value $\leq 0.5$ were retained for further analysis in step 7. |
| 3 | DNA-PAINT greyscale image rendering | The localised DNA-PAINT events are translated into a greyscale super-resolution image where pixel intensity reflects localised event density | Open DNA-PAINT localisation data with VisGUI.exe (included in the PyME suite) is used for image rendering. Use the ‘jittered triangulation’ rendering function (under the menu ‘Generate’), with default settings. | Output greyscale image must have a pixel sampling no more coarse than 5 nm/px.<br>This image may need to be cropped to include only the regions with correlative information in order to conserve memory. |
| 4. | Detection of punctate RyR densities | Discretisation of RyR puncta to enable counting and spatial statistics | Open image data with dh5view.exe (included in the PyME suite). Enable the module ‘blobFinding’ under the menu ‘Modules’. Under the object finding panel, select ‘Multi-threshold’. Key parameters include ‘Blur size’ of ~ 3.5 and ‘Threshold’ between 1.0 and 5.0. Once the detection is complete, the x-y coordinates can be saved via menu File > Save Results > Save Positions | The ‘blur size’ and ‘threshold’ parameters depend on the background intensity and the closeness of RyR clustering. The indicated parameters are optimal for RyR DNA-PAINT image data in rat ventricular cardiomyocytes. |
| 5. | Image alignment | Manual alignment of the Ca<sup>2+</sup> image against the RyR (DNA-PAINT image) to obtain the primary alignment vectors and the cross-correlation to obtain the fine-alignment vectors. | The IDL script entitled ‘align.pro’ included in the GitHub repository will perform this. Specific instructions are found in accompanying README.txt file. | Manual alignment is ideally done based on cell outline geometry. If fiducials present in the cropped image region, they can be used for confirming this alignment. In the absence of the cross-correlation step<sup>#</sup>, registration error in the manual alignment is equivalent to the resolution of the image with the lower resolution.<br><sup>#</sup> Within a 200 nm tolerance. |
| 6. | Correlation of Ca<sup>2+</sup> spark positions with RyRs | The alignment vectors are applied to appropriately align the measured x-y coordinates of the Ca<sup>2+</sup> sparks. | The IDL script entitled ‘correlative_spark.pro’ included in the GitHub repository will perform this. Specific instructions are found in accompanying README.txt file. | * Optional distortion-correction vectors could be applied to the Ca<sup>2+</sup> spark coordinates to correct for chromatic shift between imaging filter sets (see bottom of this table). |
| 7. | Interrogation of RyRs local to each Ca<sup>2+</sup> spark | RyR puncta within a circular field equivalent to the FWHM of the Ca<sup>2+</sup> spark is recorded and plotted against the ‘spark mass’ | This step is built into the IDL script entitled ‘correlative_spark.pro’ included in the GitHub repository will perform this. Specific instructions are found in accompanying README.txt file. | Spark exclusion criteria (listed in step 2 of this table) are recommended in order to reject out-of-focus sparks or spurious detections. |
| Optional (prior to imaging) | Estimating and applying chromatic shift field | To offset chromatic shift between imaging channels for Ca<sup>2+</sup> imaging and DNA-PAINT (if different filter sets and laser spots are used for the two images) | Acquire series of images of 100 nm multispectral beads in both channels. Combine the two image series vertically using FIJI menus: Image > Stacks > Tools > Combine > ‘Combine vertically’.<br>Open this combined time series stack in dh5view.exe. Run the centroid localisation protocol included within the ‘SplitterFitFNR’ module which outputs a shift field containing x and y series of vectors for the image field. (Documentation found under www.python-microscopy.org). The shift vector maps can be exported and be applied in step 6 (with the IDL code ‘correlative_spark.pro’) |  |

## 8. Discussion

### Utility of the protocol to the cell biologist

This Ca^2+^ sparks/RyR protocol, to our knowledge, provided the first opportunity for examining the structure-function relationship underpinning fast intracellular second messenger systems in wildtype primary cell-types. Many studies have previously done this comparison through second-order comparisons of Ca^2+^ spark properties and the cellular structure indirectly. A key advantage of this approach is that it is, for the first time, possible to visualise the protein organisation patterns which are *directly* responsible for the signal recorded. The spatially-constrained approach to sampling the underlying protein distribution (i.e. examining the RyR pattern which directly underlies the Ca^2+^ spark) allows the biologist to narrow their examination to the protein sub-populations which are directly responsible for a specific signal or response. In principle, this can easily be extended to examining other Ca^2+^ signalling phenotypes such as Ca^2+^ waves. For the biophysicist who is interested in the *mechanisms* underpinning the genesis of fast second-messenger signals, in silico simulations informed by non-correlative Ca^2+^ and structural image data (e.g. (4, 43–45)) have previously been the primary strategy. The experimental correlative protocol presented here allows such investigations to be extended to the living cell, for the signalling nanodomains to be pharmacologically or biophysically (e.g. electrically) perturbed to directly interrogate how protein-protein coupling and biochemical modifications (e.g. phosphorylation which alters gating kinetics) can physically re-shape the Ca^2+^ signals. More directly, this correlative protocol allows the visualisation of how multiple nanodomains or protein clusters located in the same sub-cellular neighbourhood may be recruited into producing an elementary signal (e.g. Figure 5A).

The method also benefits from the tremendous versatility and the true molecular-scale resolution offered by DNA-PAINT. RyRs, the primary mechanism of Ca^2+^ release in cardiomyocytes, typically self-assemble into clusters at a centre-to-centre distance of ~ 30-40 nm (4, 46) that allows individual channels to be resolved visually. Among additional capabilities of DNA-PAINT, we underscore ‘Quantitative PAINT’ which is an alternative way of counting targets (16), and exchange-PAINT which allows for the differential interactions of RyRs with other types of regulatory proteins locally (11).

Rapid second messenger signals such as Ca^2+^ sparks, Ca^2+^ transients and Ca^2+^ waves are cellular phenotypes which are integral to the function of excitable cell types such as neurons and muscle. Often, disease physiologies which are rooted in subtle changes in such Ca^2+^ signals (e.g. contractile dysfunction in cardiomyopathies or excitotoxicity leading to neuronal cell death) require a careful re-examination of the changes which may take place in the genesis of the signal. Protocols such as this provide a valuable experimental strategy to directly visualise *how* Ca^2+^ sparks may be fundamentally re-shaped in the contexts of disease-related remodelling of nanodomains (e.g. (4)) and/or in the presence of pharmacological agent.

### Limitations

The protocol comes with a number of limitations which can be addressed with better hardware and software. The primary limitation is the labour-intensive and time-consuming nature of the experimental pipeline of the method (i.e. steps outlined in Figure 3). There is capacity to re-purpose gridded chambers which are designed for CLEM, however we opted for the round glass dishes with the gridded substrate because of the low background fluorescence which was particularly key for achieving high localisation precision for DNA-PAINT events and the Ca^2+^ sparks. It was also necessary to maintain the cells in generally good cell health during the Ca^2+^ spark imaging which often involved imaging multiple cells within a given dish before the cells were fixed (windows up to ~ 30 mins). For highly dynamic proteins, particularly plasmalemmal proteins, such time delays could corrupt the spatially-constrained interrogation of the channels. From limited live-cell RyR imaging data available, we know that there may be some, but very limited mobility of the RyRs in the sub-plasmalemmal regions in this timescale (24). Furthermore, the use of diffusive Ca^2+^ indicator dyes worsens the measured width of the Ca^2+^ spark and can also alter the free diffusion of cytoplasmic Ca^2+^. Whilst a genetically-encoded Ca^2+^ sensor such as GCaMP6f (e.g. (47)) could vastly alleviate this limitation, it also requires more long-term culture of the cells (at least 48 hours) and spatially-uniform expression of the sensor. It is therefore limiting for examining primary cell-types.

The image acquisition and analysis steps also involve some noteworthy challenges. The use of TIRF as a selective-plane illumination allows both the Ca^2+^ and the DNA-PAINT patterns to be constrained to the sub-plasmalemmal 100-200 nm volume. We have not measured or mapped the height of the evanescent fields for the two channels of illumination, and hence assume that they are equivalent. Here, we exploited the fact that rat ventricular cardiomyocytes generally lack RyR clusters between the depths of ~ 50 nm and 600 nm from the cell surface (48). Therefore, the effect of three-dimensionality of both the Ca^2+^ signals and the RyR organisation was unlikely to corrupt the entire analysis. The distortion shift map which was used to correct any in-plane offsets between the two imaging channels did not capture any changes in axial offsets. It was assumed that TIRF illumination was always achieved within a narrow volume above the coverslip.

We found that the cell outline and shape were more useful for image alignment than the use of fluorescent beads or nanoparticles as fiducial markers, as we have demonstrated in a previous correlative tissue imaging method (39). Multi-emission beads or nanoparticles attached to the coverslip were most useful as a confirmation of the manual alignment on the basis of cell outline. In our experience, the beads immobilised on the coverslip were also useful for ensuring that both images were focused on the absolute surface of the cell. As a protocol which involves extensive solution changes (as in Figure 3) however, these particles were prone to being washed away. Where fluorescent beads were retained, we used them more as a means of validating the image alignment rather than the primary guide for it. A limitation inherent to this approach is that the Ca^2+^ image data often feature a poor signal-to-noise ratio which is reduced by taking an averaged image for delineating the cell boundary. Occasionally, smaller cell-types (e.g. fibroblasts) which remain attached to the side of the cardiomyocytes can present as an appendage with a non-zero cytoplasmic Ca^2+^ signal; however, tend to be devoid of RyR2 labelling.

For super-resolution imaging, we used DNA-PAINT in order to exploit its many benefits, including the in-plane resolution approaching 10 nm. It must be noted that this protocol is sufficiently robust to replace DNA-PAINT with more popular SMLM techniques such as dSTORM. Common to all techniques using antibody or extrinsic marker systems, is the uncertainties around incomplete target saturation, pattern distortion (depending on marker size and orientation of binding), non-specific binding and heterogeneous target or probe labelling. We used a highly-established anti-RyR antibody which has been validated previously with DNA-PAINT (11) and enhanced expansion microscopy (4) as a marker capable of resolving the majority of individual RyR channels, *in situ*. Other antibodies and targets which are combined with this protocol would ideally need to undergo similar validation.

## 9. Conclusion

We have developed an experimental and image analysis protocol which allows correlative imaging of fast, diffusive, sub-plasmalemmal second-messenger signals (Ca^2+^ sparks in the examples presented here) and true molecular-scale super-resolution imaging of the underpinning protein organisation (RyR channels) in genetically-unmodified primary cells, freshly isolated from the host. We have demonstrated how this type of data enables the only experimental strategy for a local interrogation of how individual proteins and/or clusters can be recruited to produce a Ca^2+^ spark. Ca^2+^ and other fast-diffusing second messenger signals are a key feature of excitable cells such as neurons and muscle. We therefore anticipate this protocol to enable new investigations which *directly* probe the structure-function relationship underpinning cellular physiology and human pathologies.

## 10. End Matter

### Author contributions

MEH, IJ, DS, EW & NG conceived the experiments. IJ, DS, EW, HMK, SSS, RN, EP & NG provided the supervision for the experiments. MEH carried out the experiments. MEH, TMDS, HK, SSS, ZY & RN developed the experimental materials and carried out the enzymatic cell isolations for the experiments. MEH, & IJ analysed and interpreted the data. MEH & IJ wrote the manuscript.

## Acknowledgements

The authors acknowledge Dr David Baddeley & Prof Christian Soeller for their advice on installing and adapting the PyME software for image acquisition and analysis. Further gratitude is extended to anonymous peers who have offered feedback on this manuscript to-date. The research work was funded by the Royal Society Research Grant (RG150698) awarded to IJ and the Leeds Anniversary Research Scholarship awarded to MEH.

## References

1. Sigal YM, Zhou R, Zhuang X. Visualizing and discovering cellular structures with super-resolution microscopy. Science. 2018;361(6405):880–7.

2. Greenfield D, McEvoy AL, Shroff H, Crooks GE, Wingreen NS, Betzig E, et al. Self-organization of the Escherichia coli chemotaxis network imaged with super-resolution light microscopy. PLoS Biol. 2009;7(6):e1000137.

3. Zhang J, Carver CM, Choveau FS, Shapiro MS. Clustering and Functional Coupling of Diverse Ion Channels and Signaling Proteins Revealed by Super-resolution STORM Microscopy in Neurons. Neuron. 2016;92(2):461–78.

4. Sheard TMD, Hurley ME, Colyer J, White E, Norman R, Pervolaraki E, et al. Three-Dimensional and Chemical Mapping of Intracellular Signaling Nanodomains in Health and Disease with Enhanced Expansion Microscopy. ACS Nano. 2019;13(2):2143–57.

5. Schroeder LK, Barentine AES, Merta H, Schweighofer S, Zhang Y, Baddeley D, et al. Dynamic nanoscale morphology of the ER surveyed by STED microscopy. J Cell Biol. 2019;218(1):83–96.

6. Stephan T, Roesch A, Riedel D, Jakobs S. Live-cell STED nanoscopy of mitochondrial cristae. Sci Rep. 2019;9(1):12419.

7. Vassilopoulos S, Gibaud S, Jimenez A, Caillol G, Leterrier C. Ultrastructure of the axonal periodic scaffold reveals a braid-like organization of actin rings. Nat Commun. 2019;10(1):5803.

8. Mikhaylova M, Cloin BM, Finan K, van den Berg R, Teeuw J, Kijanka MM, et al. Resolving bundled microtubules using anti-tubulin nanobodies. Nat Commun. 2015;6:7933.

9. Dudok B, Barna L, Ledri M, Szabo SI, Szabadits E, Pinter B, et al. Cell-specific STORM super-resolution imaging reveals nanoscale organization of cannabinoid signaling. Nat Neurosci. 2015;18(1):75–86.

10. Pageon SV, Tabarin T, Yamamoto Y, Ma Y, Nicovich PR, Bridgeman JS, et al. Functional role of T-cell receptor nanoclusters in signal initiation and antigen discrimination. Proc Natl Acad Sci U S A. 2016;113(37):E5454–63.

11. Jayasinghe I, Clowsley AH, Lin R, Lutz T, Harrison C, Green E, et al. True Molecular Scale Visualization of Variable Clustering Properties of Ryanodine Receptors. Cell Rep. 2018;22(2):557–67.

12. Douglass AD, Vale RD. Single-molecule microscopy reveals plasma membrane microdomains created by protein-protein networks that exclude or trap signaling molecules in T cells. Cell. 2005;121(6):937–50.

13. Jones RA, Harrison C, Eaton SL, Llavero Hurtado M, Graham LC, Alkhammash L, et al. Cellular and Molecular Anatomy of the Human Neuromuscular Junction. Cell Rep. 2017;21(9):2348–56.

14. Jungmann R, Avendano MS, Woehrstein JB, Dai M, Shih WM, Yin P. Multiplexed 3D cellular super-resolution imaging with DNA-PAINT and Exchange-PAINT. Nat Methods. 2014;11(3):313–8.

15. Jayasinghe ID, Clowsley AH, Munro M, Hou Y, Crossman DJ, Soeller C. Revealing T-Tubules in Striated Muscle with New Optical Super-Resolution Microscopy Techniquess. Eur J Transl Myol. 2015;25(1):4747.

16. Jungmann R, Avendano MS, Dai M, Woehrstein JB, Agasti SS, Feiger Z, et al. Quantitative super-resolution imaging with qPAINT. Nat Methods. 2016;13(5):439–42.

17. Clowsley AH, Kaufhold WT, Lutz T, Meletiou A, Di Michele L, Soeller C. Detecting nanoscale distribution of protein pairs by proximity dependent super-resolution microscopy. Journal of the American Chemical Society. 2020.

18. Beliveau BJ, Boettiger AN, Nir G, Bintu B, Yin P, Zhuang X, et al. In Situ Super-Resolution Imaging of Genomic DNA with OligoSTORM and OligoDNA-PAINT. Methods Mol Biol. 2017;1663:231–52.

19. Shim SH, Xia C, Zhong G, Babcock HP, Vaughan JC, Huang B, et al. Super-resolution fluorescence imaging of organelles in live cells with photoswitchable membrane probes. Proc Natl Acad Sci U S A. 2012;109(35):13978–83.

20. Jones SA, Shim SH, He J, Zhuang X. Fast, three-dimensional super-resolution imaging of live cells. Nat Methods. 2011;8(6):499–508.

21. Breedijk RMP, Wen J, Krishnaswami V, Bernas T, Manders EMM, Setlow P, et al. A live-cell super-resolution technique demonstrated by imaging germinosomes in wild-type bacterial spores. Sci Rep. 2020;10(1):5312.

22. Hartwich TMP, Hin Chung KK, Schroeder L, Bewersdorf J, Soeller C, Baddeley D. A stable, high refractive index, switching buffer for super-resolution imaging. bioRxiv. 2018:465492.

23. Rust MJ, Bates M, Zhuang X. Sub-diffraction-limit imaging by stochastic optical reconstruction microscopy (STORM). Nat Methods. 2006;3(10):793–5.

24. Hiess F, Detampel P, Nolla-Colomer C, Vallmitjana A, Ganguly A, Amrein M, et al. Dynamic and Irregular Distribution of RyR2 Clusters in the Periphery of Live Ventricular Myocytes. Biophys J. 2018;114(2):343–54.

25. Curd A, Leng J, Hughes R, Cleasby A, Rogers B, Trinh C, et al. Nanoscale pattern extraction from relative positions of sparse 3D localisations. bioRxiv; 2020.

26. Yang Z, Kirton HM, MacDougall DA, Boyle JP, Deuchars J, Frater B, et al. The Golgi apparatus is a functionally distinct Ca^*2+*^ store regulated by the PKA and Epac branches of the β_*1*_-adrenergic signaling pathway. Science Signaling. 2015;8(398):ra101.

27. Viero C, Kraushaar U, Ruppenthal S, Kaestner L, Lipp P. A primary culture system for sustained expression of a calcium sensor in preserved adult rat ventricular myocytes. Cell Calcium. 2008;43(1):59–71.

28. Ouyang K, Zheng H, Qin X, Zhang C, Yang D, Wang X, et al. Ca2+ sparks and secretion in dorsal root ganglion neurons. Proc Natl Acad Sci U S A. 2005;102(34):12259–64.

29. Cheng H, Lederer WJ, Cannell MB. Calcium sparks: elementary events underlying excitation-contraction coupling in heart muscle. Science. 1993;262(5134):740–4.

30. Cheng H, Lederer WJ. Calcium Sparks. Physiological Reviews. 2008;88(4):1491–545.

31. Sun XH, Protasi F, Takahashi M, Takeshima H, Ferguson DG, Franzini-Armstrong C. Molecular architecture of membranes involved in excitation-contraction coupling of cardiac muscle. J Cell Biol. 1995;129(3):659–71.

32. Hampton CM, Strauss JD, Ke Z, Dillard RS, Hammonds JE, Alonas E, et al. Correlated fluorescence microscopy and cryo-electron tomography of virus-infected or transfected mammalian cells. Nat Protoc. 2017;12(1):150–67.

33. Cosentino M, Canale C, Bianchini P, Diaspro A. AFM-STED correlative nanoscopy reveals a dark side in fluorescence microscopy imaging. Sci Adv. 2019;5(6):eaav8062.

34. Hirvonen LM, Cox S. STORM without enzymatic oxygen scavenging for correlative atomic force and fluorescence superresolution microscopy. Methods and Applications in Fluorescence. 2018;6(4):045002.

35. Gomez-Varela AI, Stamov DR, Miranda A, Alves R, Barata-Antunes C, Dambournet D, et al. Simultaneous co-localized super-resolution fluorescence microscopy and atomic force microscopy: combined SIM and AFM platform for the life sciences. Sci Rep. 2020;10(1):1122.

36. Bondia P, Casado S, Flors C. Correlative Super-Resolution Fluorescence Imaging and Atomic Force Microscopy for the Characterization of Biological Samples. Methods Mol Biol. 2017;1663:105–13.

37. Hamel V, Guichard P, Fournier M, Guiet R, Fluckiger I, Seitz A, et al. Correlative multicolor 3D SIM and STORM microscopy. Biomed Opt Express. 2014;5(10):3326–36.

38. Dudok B, Barna L, Ledri M, Szabó SI, Szabadits E, Pintér B, et al. Cell-specific STORM super-resolution imaging reveals nanoscale organization of cannabinoid signaling. Nature neuroscience. 2015;18(1):75–86.

39. Crossman DJ, Hou Y, Jayasinghe I, Baddeley D, Soeller C. Combining confocal and single molecule localisation microscopy: A correlative approach to multi-scale tissue imaging. Methods. 2015;88:98–108.

40. Gwathmey JK, Hajjar RJ, Solaro RJ. Contractile deactivation and uncoupling of crossbridges. Effects of 2,3-butanedione monoxime on mammalian myocardium. Circ Res. 1991;69(5):1280–92.

41. Steele EM, Steele DS. Automated detection and analysis of Ca(2+) sparks in x-y image stacks using a thresholding algorithm implemented within the open-source image analysis platform ImageJ. Biophysical journal. 2014;106(3):566–76.

42. Baddeley D, Cannell MB, Soeller C. Visualization of localization microscopy data. Microsc Microanal. 2010;16(1):64–72.

43. Hollingworth S, Peet J, Chandler WK, Baylor SM. Calcium sparks in intact skeletal muscle fibers of the frog. J Gen Physiol. 2001;118(6):653–78.

44. Colman MA, Pinali C, Trafford AW, Zhang H, Kitmitto A. A computational model of spatio-temporal cardiac intracellular calcium handling with realistic structure and spatial flux distribution from sarcoplasmic reticulum and t-tubule reconstructions. PLoS Comput Biol. 2017;13(8):e1005714.

45. Rajagopal V, Bass G, Walker CG, Crossman DJ, Petzer A, Hickey A, et al. Examination of the Effects of Heterogeneous Organization of RyR Clusters, Myofibrils and Mitochondria on Ca2+ Release Patterns in Cardiomyocytes. PLoS Comput Biol. 2015;11(9):e1004417.

46. Walker MA, Kohl T, Lehnart SE, Greenstein JL, Lederer WJ, Winslow RL. On the Adjacency Matrix of RyR2 Cluster Structures. PLoS Comput Biol. 2015;11(11):e1004521.

47. Yin CC, Lai FA. Intrinsic lattice formation by the ryanodine receptor calcium-release channel. Nat Cell Biol. 2000;2(9):669–71.

48. Shang W, Lu F, Sun T, Xu J, Li LL, Wang Y, et al. Imaging Ca2+ nanosparks in heart with a new targeted biosensor. Circ Res. 2014;114(3):412–20.

49. Hou Y, Jayasinghe I, Crossman DJ, Baddeley D, Soeller C. Nanoscale analysis of ryanodine receptor clusters in dyadic couplings of rat cardiac myocytes. J Mol Cell Cardiol. 2015;80:45–55.

